# Antimicrobial peptides and proteins as alternative antibiotics for porcine semen preservation

**DOI:** 10.1101/2023.10.31.564986

**Authors:** Jose Luis Ros-Santaella, Pavel Nový, Maria Scaringi, Eliana Pintus

**Affiliations:** Department of Veterinary Sciences, Faculty of Agrobiology, Food, and Natural Resour+ces, Czech University of Life Sciences Prague, Kamýcká 129, 165 00 Praha, 6-Suchdol, Czech Republic; Department of Food Science. Faculty of Agrobiology, Food, and Natural Resour+ces, Czech University of Life Sciences Prague, Kamýcká 129, 165 00 Praha, 6-Suchdol, Czech Republic

## Abstract

**Background:** Antimicrobial resistance (AMR) is nowadays a major emerging challenge for public health worldwide. The over- and misuse of antibiotics, including those for cell culture, are promoting AMR while also encouraging the research and employment of alternative drugs. The addition of antibiotics to the cell media is a must in sperm preservation, being gentamicin the most used for boar semen. Because of its continued use, several bacterial strains present in boar semen have developed resistance to this antibiotic. Antimicrobial peptides and proteins (AMPPs) are promising candidates as alternative antibiotics because their mechanism of action is less likely to promote AMR. In the present study, we tested two AMPPs (lysozyme and nisin; 50 and 500 µg/mL) as possible substitutes of gentamicin for boar semen preservation up to 48 h of storage.

**Results:** We found that both AMPPs improved sperm plasma membrane and acrosome integrity during semen storage. The highest concentration tested for lysozyme also kept the remaining sperm parameters unaltered, at 48 h of semen storage, and reduced the bacterial load at comparable levels of the samples supplemented with gentamicin (p>0.05). On the other hand, while nisin (500 µg/mL) reduced the total Enterobacteriaceae counts, it also decreased the rapid and progressive sperm population and the seminal oxidation-reduction potential (p<0.05).

**Conclusions:** The protective effect of lysozyme on sperm function together with its antimicrobial activity and inborn presence in body fluids, including semen and cervical mucus, makes this enzyme a promising antimicrobial agent for boar semen preservation.

## Introduction

Antimicrobial resistance (AMR) is nowadays one of the main global health threats that increases the risk of disease spread, severe illness, and death. In bacteria, where AMR naturally occurs, the misuse or overuse of antibiotics are accelerating the process jeopardizing the success of modern medicine in treating infections [1].

Sperm preservation also promotes the emergence of superbugs resistant to the most common antibiotics used for this purpose [2]. Pig breeding is mainly carried out by artificial insemination (AI) using refrigerated diluted semen and entails the collection, processing, and preservation of male gametes. Even with the highest hygienic standards, bacterial contamination frequently occurs during the semen collection and handling process [3]. In addition, the liquid-storage of boar semen at 15-17 °C also favours microbes’ proliferation. All these steps make the addition of antibiotics to the semen samples a must. However, AMR to the most common antibiotics added to the semen extenders promotes the contamination of the seminal doses ranging from 14.73% to 32% [4, 5] where Gram-negative (G-) bacteria are predominant. On the other hand, most antibiotics exert negative effects *per se* (cytotoxicity) on sperm physiology [6, 7] and promote the release of bacterial endotoxins, such as lipopolysaccharide (LPS), which seriously compromises sperm function [8, 9].

The use of alternative antimicrobial agents that delay or avoid AMR is therefore urgently needed. This kind of compounds includes antimicrobial peptides and proteins (AMPPs), which are a diverse group of molecules produced by a wide variety of organisms (prokaryotes and eukaryotes) as their first line of defence against pathogenic microorganisms [10]. While the great part of these AMPPs can directly kill a wide variety of microbial pathogens (e.g., bacteria, yeasts, fungi, viruses, etc.), others modulate the host immunity [11, 12]. However, many AMPPs have a limited spectrum of activity and are effective only at very high concentrations [13] and thus, increasing their cytotoxicity if any. However, through the application of different stimuli (e.g., heat treatment) and combination with other compounds (e.g., chelators, dextran), many AMPPs can broaden their effectiveness of killing/inhibiting both Gram-positive and Gram-negative bacteria and reduce the dosage needed for that purpose [13, 14].

Nisin, a polypeptide bacteriocin produced by *Lactococcus lactis* subsp. *lactis*, is widely used (over 50 countries) as a food preservative (E-234) of meat and dairy products and a promising compound for biomedical applications such as alternative antimicrobial and cancer therapeutic [15, 16]. On the other hand, lysozyme or muramidase was the first discovered antimicrobial protein (enzyme) by Alexander Fleming [17]. It is typically found in body fluids (e.g., saliva, tears, milk, semen, cervical mucus), organs, tissues, and cells (e.g., polymorphonuclear leukocytes) from many organisms mainly acting as an inborn immunological defence against a wide variety of pathogens [18–20]. The main antibacterial spectrum of both nisin and lysozyme is Gram-positive (G+) bacteria, but, in presence of chelating agents, like ethylenediaminetetraacetic acid (EDTA), they are also effective against G-bacteria [13]. The addition of EDTA to the semen extenders is a common praxis to block the action of calcium as a mediator of sperm capacitation and acrosome reaction [21], while it could be also harnessed to increase the antimicrobial spectrum of AMPPs. For this reason, semen extenders containing chelating agents such as EDTA could be suitable in the fight against AMR using alternative antibiotics for semen preservation.

The main goal of this study was to evaluate the potential use of nisin and lysozyme as alternative antibiotics for the preservation of boar semen at 17 °C. For this purpose, we used Beltsville Thawing Solution (BTS) as semen extender that contains EDTA and sodium bicarbonate which have shown their own antimicrobial activity [22, 23] and broaden the spectrum of these AMPPs also to G-bacteria. This short-term extender allows the preservation of sperm cells for 1-3 days, although after 48 h of storage, there is a decline in the fertility rates and number of piglets born [21]. For this reason and because 85% of the AIs are carried out within the first two days after semen collection [24], the analyses of the present study were performed till 48 h of semen storage. The presence of lysozyme in several mammalian body fluids and the approval of nisin as a safe food preservative make these AMPPs suitable alternative antibiotics for boar semen preservation.

## Material and methods

All reagents were purchased from Merck (Prague, Czech Republic), unless otherwise indicated.

### Semen collection and processing

Semen was collected by the gloved hand method from fertile boars (Duroc breed) at a pig breeding company (Lipra Pork, a.s., Czech Republic). The semen from 12 boars was used in this study. Twenty mL of each ejaculate was transported to the laboratory in sterilized tubes. An aliquot of each sample was placed in 0.3% formaldehyde in phosphate-buffered saline (PBS) for assessing sperm abnormalities (200 sperm evaluated per sample). The ejaculates with more than 25% of sperm abnormalities were discarded for the experiments. Then, all ejaculates were supplemented with 5 mL of BTS (D-glucose 37 g/L, sodium citrate 6 g/L, ethylenediaminetetraacetic acid 1.25 g/L, sodium bicarbonate 1.25 g/L, potassium chloride 0.75 g/L) without antibiotic. The BTS’ pH (adjusted with NaOH 10 M) and osmolality were ∼ 7.2 (Five Easy F20, Mettler-Toledo, Switzerland) and ∼ 330 mOsm/kg H_2_O (Osmo-mat 3000, Gonotec, Germany), respectively. The BTS was prepared, under sterile conditions, in water (Carl Roth) and filtered (0.2 µm filter pore, Whatman plc, United Kingdom) after preparation. For each experimental replicate, semen from three boars was pooled and centrifuged at 167 *g* for 3 min at 17 °C to remove debris and abnormal cells [25]. A sub-sample of the supernatant (fixed in 0.3% formaldehyde in PBS) was collected for the assessment of sperm concentration in a Bürker chamber. The pooled semen was then diluted to 40×10^6^ /mL in BTS extender w/o antibiotic. Four replicates were used in the present study. The initial sperm motility in all replicates was >75%.

### Treatments

A stock solution (2 mg/mL) of both nisin (N5764; ≥900 IU/mg) and lysozyme (1052810001; from egg white; ≥30 000 FIP-U/mg) was freshly prepared in BTS. Then, the solutions were filtered by a syringe filter (0.2 µm pore; Whatman plc, United Kingdom) and placed in sterile tubes. The diluted semen was split into 6 sterile tubes (diluted down 20×10^6^ spermatozoa/mL in BTS w/ or w/o gentamicin or AMPPs) to reach the final concentrations as it follows: Ctr (BTS w/o antibiotic), Gent (gentamicin, 250 µg/mL), Lys500 (lysozyme, 500 µg/mL), Lys50 (lysozyme, 50 µg/mL), Nis500 (nisin, 500 µg/mL), and Nis50 (nisin, 50 µg/mL). The choice of the used AMPPs concentration was based on their antimicrobial activity in combination with EDTA previously reported [13].

### Sperm motility and kinetics parameters

A sperm aliquot (2 µL) was loaded into a pre-warmed Leja chamber (Leja Products BV, The Netherlands, chamber depth: 20 µm). The sperm motility and kinetic parameters were evaluated, as previously reported [26], using a Computer Assisted Sperm Analysis (CASA) (NIS-Elements, Nikon, Tokyo, Japan and Laboratory Imaging, Prague, Czech Republic), which consists of an Eclipse E600 tri-ocular phase contrast microscope (Nikon, Tokyo, Japan), equipped with a warming stage set at 38 °C (Tokai Hit, Shizuoka, Japan), and a DMK 23UM021 digital camera (The Imaging Source, Bremen, Germany). The analysis was carried out using a 10× negative phase-contrast objective (Nikon, Tokyo, Japan). A total of eight descriptors of sperm motility parameters were recorded: total motility (TM, %), progressive motility (PM, %), average path velocity (VAP, µm/s), curvilinear velocity (VCL, µm/s), straight-line velocity (VSL, µm/s), amplitude of lateral head displacement (ALH, µm), beat-cross frequency (BCF, Hz), linearity (LIN, %), and straightness (STR, %). The standard CASA settings were as follows: frames per second, 60; minimum of frames acquired, 31; number of fields analysed, 6; VAP ≥ 10 µm/s to classify a spermatozoon as motile; STR ≥ 80% to classify a spermatozoon as progressive. A minimum of 200 motile sperm cells were analysed per sample. All videos were visually checked by the same researcher to remove debris or erroneously crossed sperm tracks. Sperm motile subpopulations were determined on the whole sperm population by cluster analysis.

### Sperm plasma membrane integrity, acrosomal status, and mitochondrial activity

Sperm analyses were carried out as previously described [27]. Briefly, for the assessment of membrane integrity, the sperm samples were incubated with propidium iodide (stock solution: 0.5 mg/mL in phosphate-buffered saline, PBS), carboxyfluorescein diacetate (stock solution: 0.46 mg/mL in dimethyl sulfoxide, DMSO), and formaldehyde solution (0.3%) for 10 min at 38 °C in the dark. Then, the spermatozoa were assessed under epi-fluorescence microscopy (Nikon Eclipse E600, Nikon, Japan; 40× objective), and those with green fluorescence over the entire head were considered to have an intact plasma membrane. For the acrosomal status, the percentage of sperm with a normal apical ridge (NAR) was determined. The sperm samples were fixed in a glutaraldehyde solution (2%) and evaluated under phase contrast microscopy (40× objective). To determine mitochondrial activity, the aliquots of the sperm samples were incubated with rhodamine 123 (5 mg/mL in DMSO) and propidium iodide (0.5 mg/mL in PBS) for 15 min at 38 °C in the dark. After that, the samples were centrifuged at 500 g for 5 min, the supernatant was removed, and the sperm pellet was resuspended in PBS. Then, the spermatozoa were evaluated using epi-fluorescence microscopy (40× objective), and the spermatozoa showing a bright green fluorescence over the midpiece were considered to have a high mitochondrial activity. Two-hundred sperm cells were assessed per analysis by the same observer.

### Seminal oxidation-reduction potential (ORP)

The seminal ORP of the samples were determined as previously reported [26] with minor modifications. At the end of sperm incubation, a sample from each treatment was centrifuged at 16, 300 *g* for 5 minutes at room temperature. Then, 800 μL of the supernatant was transferred into a microcentrifuge tube and incubated at 38 °C. The ORP was measured using a micro ORP electrode with Argenthal™ reference system and platinum ring (InLab® Redox Micro, Mettler-Toledo, Switzerland) connected to a pH/mV meter (Five Easy F20, Mettler-Toledo, Switzerland). The ORP of each sample was recorded after embedding the microelectrode into the solution for 3 minutes. After each sample analysis, the probe was calibrated into a redox buffer solution (220 mV, pH 7, Mettler-Toledo, Switzerland) for 30 seconds. The assay was run in duplicate per each sample and expressed in millivolts (mV). The ORP levels were not normalized [28], because the experiments were performed at the same sperm concentration (i.e., 20×10^6^/mL).

### Isolation of contaminating bacteria and MALDI-TOF MS identification

Pseudomonas (PA) agar (Sigma-Aldrich, Prague, CZ), Blood (BA), MacConkey (MCA), Mannitol Salt (MSA), Plate count (PCA) agars (Oxoid, Basigstoke, UK) were used for the isolation of contaminating bacteria. Sample aliquots of 50 µl were plated in duplicate on 90 mm agar plates using spiral plate inoculator EasySpiral (Interscience, Saint Nom, France) and incubated aerobically at 37°C for 24–48 h. Selected colonies with different morphology were further repassaged to ensure a pure culture.

Freshly grown colonies were harvested and subjected to the standard procedure recommended by Bruker Daltonics for the MALDI-TOF MS identification (ethanol-formic acid extraction procedure and then mixed with HCCA matrix). Protein spectra were measured and processed by Autoflex Speed MALDI-TOF MS using FlexControl 3.4; MALDI Biotyper Compass version 4.1; and flexAnalysis version 3.4 software (all Bruker Daltonics, Bremen, Germany).

### Bacterial counts determination

The bacterial contamination rates in pooled boar ejaculates in the BTS extender were determined the first day following the insemination doses preparation (Ctr), and then after 24 and 48 h of semen storage at 17°C for all treatments. The PCA and MCA agars were used for the enumeration of total aerobic mesophilic bacteria and Enterobacteriaceae, respectively. Sample aliquots of 100 µl were plated in duplicate using a spread plate technique. Samples diluted 10-fold in sterile peptone saline were plated in duplicate using a spiral plate inoculator with a 10^-5^ dilution rate. The inoculated plates were incubated for 24–48 h at 37°C. The microbial counts obtained by the spiral plate technique were interpreted according to the NF V08-100 Standard [29]. The final counts were expressed as log CFU/mL (CFU: colony forming unit) of an insemination dose. Because of the volume used for the initial microbial culture (100 µl; undiluted samples), a value of 0 in the bacterial counts is equivalent to <10 CFU/mL.

### Statistical analyses

The statistical analyses were carried out by the SPSS 24 statistical software package (IBM Inc, Chicago, IL, USA). To determine sperm motile subpopulations, a two-step cluster analysis was applied to the whole sperm population using VAP and STR as variables. The number of clusters was automatically determined using the Euclidean distance measure and the Schwarz’s Bayesian criterion. After that, the number of clusters previously obtained was used to set up the K-means cluster analysis by using the iteration and classification method. The Kruskal-Wallis test was used to check for differences between sperm motile subpopulations in kinetic variables. A generalized linear model was used to analyse the effects of the treatments and storage times on the sperm variables and bacterial counts. The data concerning the bacterial load were log-transformed to perform the analyses. The data are expressed as the mean ± standard error. The statistical significance was set at p<0.05.

## Results

### Sperm motility and kinetic parameters

The average values of motility and kinetics of boar spermatozoa during semen storage are shown in Table 1. At 24 h, there were no significant differences between Gent and Ctr groups in the total motility (p>0.05). Interestingly, Lys50 and Nis50 showed greater total motility than the Gent group (p<0.05). On the other hand, a significant (p<0.05) decrease in progressive motility was observed in Nis treatments in comparison with Gent group. Overall, most kinetic parameters (VAP, VCL, VSL, and ALH) of the Gent group were higher (p<0.05) than those of other treatments (Ctr included). On the other hand, Lys and Nis treatments (500 µg/mL) showed higher values of VSL (Lys only), BCF (both treatments), and LIN (Lys only) than the Ctr group (p<0.05).

**Table 1.** Effect of lysozyme and nisin on sperm motility and kinetic parameters during porcine semen storage at 17 °C.

| Time | Treatment | TM (%) | PM (%) | VAP (μm/s) | VCL (μm/s) | VSL (μm/s) | ALH (μm) | BCF (Hz) | LIN (%) | STR (%) |
| --- | --- | --- | --- | --- | --- | --- | --- | --- | --- | --- |
| **0 h** | Gent | 79.67±3.72 | 52.75±1.49 | 82.55±5.87 | 132.95±9.90 | 65.51±3.90 | 5.31±0.28 | 15.87±0.45 | 50.01±1.07 | 77.90±0.92 |
| **24 h** | Gent | 63.25±1.78<sup>bc</sup> | 56.27±3.49<sup>a</sup> | 99.44±1.25<sup>a</sup> | 146.65±1.11<sup>a</sup> | 86.43±2.33<sup>a</sup> | 5.64±0.08<sup>a</sup> | 17.39±0.31<sup>ab</sup> | 59.31±0.99<sup>a</sup> | 85.75±1.01<sup>a</sup> |
|  | Ctr | 68.13±4.11<sup>ac</sup> | 49.36±4.64<sup>abc</sup> | **83.17±1.51<sup>b</sup>** | **126.80±4.23<sup>b</sup>** | **65.59±2.80<sup>bd</sup>** | **4.97±0.17<sup>b</sup>** | 16.85±0.29<sup>b</sup> | **53.24±3.53<sup>bc</sup>** | **78.41±3.79<sup>bc</sup>** |
|  | Lys500 | 65.03±2.61<sup>bcd</sup> | 53.44±3.56<sup>ab</sup> | **87.15±2.48<sup>b</sup>** | **126.41±4.13<sup>b</sup>** | **73.81±1.91<sup>c</sup>** | **5.03±0.21<sup>b</sup>** | 17.79±0.41<sup>a</sup> | 59.48±2.00<sup>a</sup> | 83.99±2.21<sup>ab</sup> |
|  | Lys50 | **72.51±2.09<sup>a</sup>** | 52.35±2.64<sup>abc</sup> | **82.80±1.95<sup>b</sup>** | **123.98±2.99<sup>b</sup>** | **67.14±2.58<sup>b</sup>** | **4.83±0.11<sup>b</sup>** | 17.19±0.40<sup>ab</sup> | 54.74±2.00<sup>ab</sup> | **80.01±1.97<sup>bc</sup>** |
|  | Nis500 | 60.17±4.48<sup>b</sup> | **47.51±1.89<sup>bc</sup>** | **84.47±1.85<sup>b</sup>** | **123.21±3.90<sup>b</sup>** | **68.50±1.04<sup>bc</sup>** | **4.94±0.16<sup>b</sup>** | 17.68±0.38<sup>a</sup> | 57.00±1.35<sup>ac</sup> | **79.93±1.32<sup>bc</sup>** |
|  | Nis50 | **69.61±1.26<sup>ad</sup>** | **45.72±4.96<sup>c</sup>** | **81.07±2.85<sup>b</sup>** | **123.47±5.79<sup>b</sup>** | **60.84±3.55<sup>d</sup>** | **4.76±0.17<sup>b</sup>** | 17.02±0.22<sup>ab</sup> | **50.78±3.81<sup>b</sup>** | **74.94±4.30<sup>c</sup>** |
| **48 h** | Gent | 63.50±1.27<sup>ab</sup> | 53.25±1.20 | 91.16±2.56<sup>a</sup> | 153.37±4.24<sup>a</sup> | 78.22±2.29<sup>a</sup> | 5.28±0.12<sup>ab</sup> | 16.10±0.26<sup>ab</sup> | 51.41±0.55<sup>ab</sup> | 84.44±1.18<sup>a</sup> |
|  | Ctr | 63.65±2.19<sup>ab</sup> | 47.19±4.78 | **84.56±2.41<sup>bc</sup>** | 146.67±7.39<sup>ab</sup> | **67.84±0.75<sup>bd</sup>** | 5.21±0.16<sup>ab</sup> | 15.50±0.29<sup>b</sup> | 47.24±2.67<sup>b</sup> | 78.87±3.06<sup>ab</sup> |
|  | Lys500 | 65.79±0.96<sup>ab</sup> | 54.21±2.49 | 87.93±0.82<sup>ab</sup> | 143.62±2.17<sup>ab</sup> | 75.25±2.24<sup>ac</sup> | 5.24±0.05<sup>ab</sup> | 16.55±0.10<sup>a</sup> | 52.89±1.93<sup>a</sup> | 83.97±1.83<sup>ab</sup> |
|  | Lys50 | 67.70±2.48<sup>a</sup> | 49.24±1.90 | 88.86±0.87<sup>ab</sup> | 151.50±1.67<sup>ab</sup> | 72.61±1.71<sup>ab</sup> | 5.43±0.10<sup>a</sup> | 15.66±0.21<sup>b</sup> | 48.53±1.11<sup>ab</sup> | 80.23±1.77<sup>ab</sup> |
|  | Nis500 | 60.67±1.30<sup>b</sup> | 46.99±2.26 | **83.82±3.26<sup>bc</sup>** | **138.96±6.75<sup>b</sup>** | **69.66±3.96<sup>bcd</sup>** | 5.08±0.27<sup>ab</sup> | 16.02±0.30<sup>ab</sup> | 51.41±2.09<sup>ab</sup> | 82.07±2.29<sup>ab</sup> |
|  | Nis50 | 68.48±2.83<sup>a</sup> | 49.16±2.57 | **81.66±2.09<sup>c</sup>** | **140.20±7.19<sup>b</sup>** | **64.91±1.38<sup>d</sup>** | 4.94±0.25<sup>b</sup> | 15.61±0.43<sup>b</sup> | 47.38±2.37<sup>b</sup> | **78.76±2.26<sup>b</sup>** |

Different superscript letters indicate significant differences (p<0.05) between treatments within each given time. Bold numbers indicate significant differences (p<0.05) between treatments (including Ctr) and Gent group. Gent = gentamicin; Ctr = control; Lys = lysozyme 500; Nis = Nisin; Treatments = Lys500 (500 μg/mL); Lys50 (50 μg/mL); Nis500 (500 μg/mL); Nis50 (50 μg/mL). TM = total motility; PM = progressive motility; VAP = average path velocity; VCL = curvilinear velocity; VSL = straight-line velocity; ALH = amplitude of lateral head displacement; BFC = beat-cross frequency; LIN = linearity; STR = straightness. The data are shown as the mean ± standard error of four replicates.

At 48 h, Lys treatments did not differ (p>0.05) in any of the motility and kinetic parameters when compared with the Gent group. By contrast, Ctr and Nis treatments showed lower values than the Gent group in several kinetic parameters (p<0.05). Similarly to the results observed at 24 h, Lys500 showed higher values of VSL, BCF, and LIN than the Ctr group (p<0.05).

The motility subpopulation analyses rendered three groups (Table 2) as follows: SP1 (rapid and progressive spermatozoa), SP2 (rapid and no progressive spermatozoa), and SP3 (slow and no progressive spermatozoa). The percentage of each subpopulation for every treatment and during semen storage is shown in Table 3. Interestingly, there were no significant differences (p>0.05) between Gent and Lys treatments in any of the sperm subpopulations during semen storage. In addition, Lys treatments showed more rapid and progressive sperm (SP1) than the Ctr group (p<0.05). On the other hand, Ctr and Nis treatments showed a higher percentage of non-progressive spermatozoa (SP2 and SP3) when compared to the Gent and Lys treatments (p<0.05).

**Table 2.** Sperm subpopulations based on kinetic parameters during porcine semen storage at 17 °C.

| Sperm subpopulations | Number of spermatozoa | VAP (µm/s) | VCL (µm/s) | VSL (µm/s) | ALH (µm) | BCF (Hz) | LIN (%) | STR (%) |
| --- | --- | --- | --- | --- | --- | --- | --- | --- |
| **Rapid and progressive spermatozoa (SP1)** | 13,095<br>(49.21%) | 92.64±0.25<sup>a</sup> | 143.35±0.41<sup>b</sup> | 85.00±0.26<sup>a</sup> | 5.40±0.02<sup>a</sup> | 17.10±0.06<sup>a</sup> | 60.61±0.12<sup>a</sup> | 91.23±0.07<sup>a</sup> |
| **Rapid and no progressive spermatozoa (SP2)** | 2,755<br>(10.35%) | 89.17±0.45<sup>b</sup> | 153.90±0.88<sup>a</sup> | 32.10±0.33<sup>b</sup> | 5.25±0.03<sup>a</sup> | 16.14±0.09<sup>a</sup> | 22.27±0.24<sup>b</sup> | 36.60±0.35<sup>b</sup> |
| **Slow and no progressive spermatozoa (SP3)** | 10,760<br>(40.44%) | 6.76±0.10<sup>c</sup> | 16.47±0.21<sup>c</sup> | 4.36±0.08<sup>c</sup> | 0.75±0.01<sup>b</sup> | 9.29±0.04<sup>b</sup> | 21.04±0.16<sup>c</sup> | 52.38±0.25<sup>c</sup> |

Different superscript letters indicate significant differences ( $p < 0.05$ ) in kinetic parameters between the different subpopulations. VAP = average path velocity; VCL = curvilinear velocity; VSL = straight-line velocity; ALH = amplitude of lateral head displacement; BFC = beat-cross frequency; LIN = linearity; STR = straightness. The data are shown as the mean ± standard error of four replicates.

**Table 3.** Effect of lysozyme and nisin on sperm subpopulations based on kinetic parameters during porcine semen storage at 17 °C.

| Time | Treatment | Rapid and progressive spermatozoa (SP1, %) | Rapid and no progressive spermatozoa (SP2, %) | Slow and no progressive spermatozoa (SP3, %) |
| --- | --- | --- | --- | --- |
| **24 h** | Gent | 53.16±2.00<sup>ac</sup> | 7.13±1.00<sup>b</sup> | 39.71±1.72<sup>bc</sup> |
|  | Ctr | 47.98±1.77<sup>bc</sup> | **13.84±4.66<sup>a</sup>** | 38.18±4.26<sup>bc</sup> |
|  | Lys500 | 51.42±1.40<sup>ac</sup> | 7.97±2.50<sup>b</sup> | 40.61±2.87<sup>b</sup> |
|  | Lys50 | 53.97±2.63<sup>a</sup> | 11.92±2.11<sup>ab</sup> | 34.11±2.14<sup>c</sup> |
|  | Nis500 | **43.18±3.35<sup>b</sup>** | 10.08±1.20<sup>ab</sup> | **46.74±3.91<sup>a</sup>** |
|  | Nis50 | **44.91±4.36<sup>b</sup>** | **15.63±3.98<sup>a</sup>** | 39.46±2.23<sup>bc</sup> |
| **48 h** | Gent | 51.40±0.77<sup>ac</sup> | 7.99±1.18 | 40.60±1.71<sup>ab</sup> |
|  | Ctr | 46.92±2.73<sup>bc</sup> | 11.09±2.4 | 42.00±2.98<sup>ab</sup> |
|  | Lys500 | 52.87±1.31<sup>a</sup> | 7.94±1.28 | 39.18±0.87<sup>b</sup> |
|  | Lys50 | 51.34±1.49<sup>ac</sup> | 12.00±2.17 | 36.66±2.72<sup>b</sup> |
|  | Nis500 | **45.29±2.17<sup>b</sup>** | 8.53±1.91 | 46.18±0.93<sup>a</sup> |
|  | Nis50 | 49.09±2.17<sup>ab</sup> | 11.27±1.65 | 39.64±1.65<sup>b</sup> |

Different superscript letters indicate significant differences ( $p < 0.05$ ) between treatments within each given time. Bold numbers indicate significant differences ( $p < 0.05$ ) between treatments (including Ctr) and Gent group. Gent = gentamicin; Ctr = control; Lys = lysozyme 500; Nis = Nisin; Treatments = Lys500 (500 µg/mL); Lys50 (50 µg/mL); Nis500 (500 µg/mL); Nis50 (50 µg/mL). The data are shown as the mean ± standard error of four replicates.

### Sperm plasma membrane integrity, acrosomal status, and mitochondrial activity

The results are shown in Figure 1. At 24 h, the percentage of sperm with intact plasma membrane did not differ between treatments (p>0.05). At the same incubation time, there were no differences between Gent and Ctr groups in the acrosomal status (p>0.05). Interestingly, Lys500 better preserved the acrosome integrity (p<0.05) when compared to Gent, while the remaining Lys and Nis treatments did not show any significant differences (p>0.05) with the Gent group. Both Lys500 and Nis treatments showed a higher percentage of sperm with intact acrosome than the Ctr group (p<0.01). The mitochondrial activity did not show any significant difference between treatments (p>0.05).

**Figure 1.**
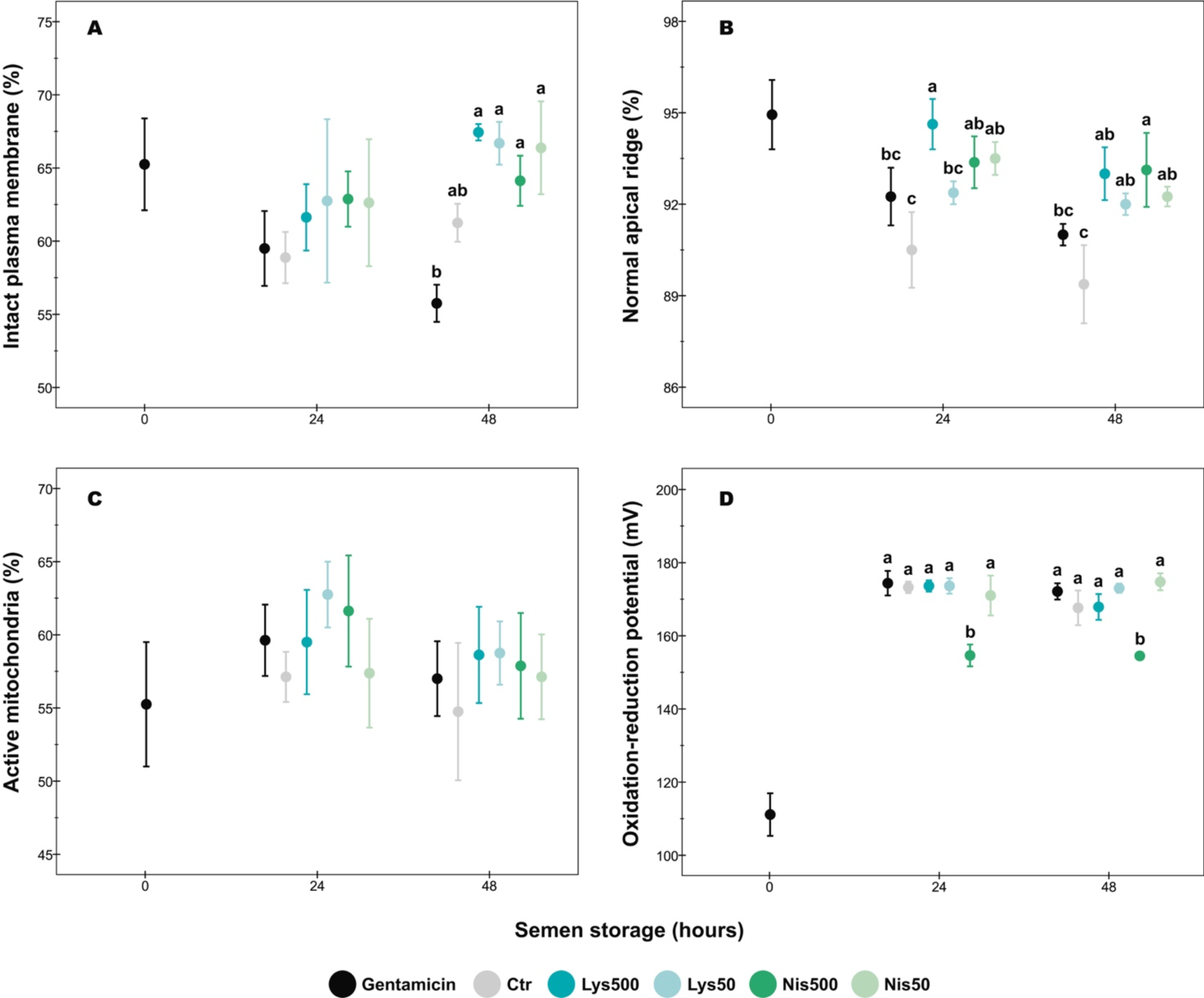
Effect of lysozyme and nisin on sperm parameters during porcine semen storage at 17 °C. A) Sperm plasma membrane integrity; B) Sperm acrosomal status; C) Sperm mitochondrial activity; D) Seminal oxidation-reduction potential. Different superscript letters indicate significant differences (p<0.05) between treatments within each given time. The data are shown as the mean ± standard error of four replicates.

At 48 h, all Lys and Nis treatments showed a higher percentage of sperm with an intact plasma membrane when compared to the Gent group (p≤0.01), whereas there were no differences between the Gent and Ctr groups (p>0.05). Regarding the acrosomal status, both Gent and Ctr groups showed similar results (p>0.05), while all Lys and Nis treatments better preserved the integrity of this organelle in comparison with the Ctr group (p≤0.02). Moreover, Nis500 better preserved the acrosome integrity than Gent (p<0.05). No significant differences were detected between groups in the mitochondrial activity (p>0.05).

### Seminal oxidation-reduction potential (ORP)

The seminal ORP was significantly lower in Nis500 treatment than in other groups (Figure 1-D; p≤0.001) at both incubation times. The remaining groups did not show any significant differences between them at any incubation period (p>0.05).

### Bacteriological profile

The bacteriological profile of boar semen samples is shown in Table 4. A total of 17 species belonging to 11 bacteria genera were identified. Gram-negative bacteria were the most prevalent contaminants in terms of frequency and the number of isolated genera (7/11). Thus, *Pseudomonas aeruginosa* (100%)*, Stenotrophomonas maltophilia* (75%), and *Klebsiella aerogenes* (50%) were the most frequent isolated species. On the other hand, *Staphylococcus* spp. were present in all the replicates as the most recurrent G+ bacteria with a total of six species isolated.

**Table 4.** Bacteriological profile of diluted porcine semen samples.

| Species | Gram classification | Replicate |  |  |  | Frequency per sample (%) |
| --- | --- | --- | --- | --- | --- | --- |
|  |  | I. | II. | III. | IV. |  |
| *Achromobacter* sp. | G- | + |  |  |  | 25 |
| *Bacillus* sp. | G+ | + |  |  |  | 25 |
| *Citrobacter koseri* | G- |  |  |  | + | 25 |
| *Corynebacterium* sp. | G+ | + |  |  |  | 25 |
| *Enterococcus faecalis* | G+ | + |  |  |  | 25 |
| *Escherichia coli* | G- |  |  | + |  | 25 |
| *Klebsiella aerogenes* | G- |  | + |  | + | 50 |
| *Klebsiella pneumoniae* | G- |  | + |  |  | 25 |
| *Pasteurella aerogenes* | G- |  |  | + |  | 25 |
| *Pseudomonas aeruginosa* | G- | + | + | + | + | 100 |
| *Staphylococcus chromogenes* | G+ | + |  | + |  | 50 |
| *Staphylococcus epidermidis* | G+ |  |  | + | + | 50 |
| *Staphylococcus hominis* | G+ |  | + |  |  | 25 |
| *Staphylococcus pasteurii* | G+ | + |  | + |  | 50 |
| *Staphylococcus pettenkoferi* | G+ |  | + |  |  | 25 |
| *Staphylococcus warneri* | G+ |  | + |  |  | 25 |
| *Stenotrophomonas maltophilia* | G- | + | + |  | + | 75 |

Sp.=species; G-=Gram-negative; G+=Gram-positive

### Total bacterial and Enterobacteriaceae counts in the semen samples (TBC and TEC)

The data relative to bacterial load of the samples are shown in Figure 2. We did not observe any bacterial growth (TBC and TEC) in the Gent group during semen storage.

**Figure 2.**
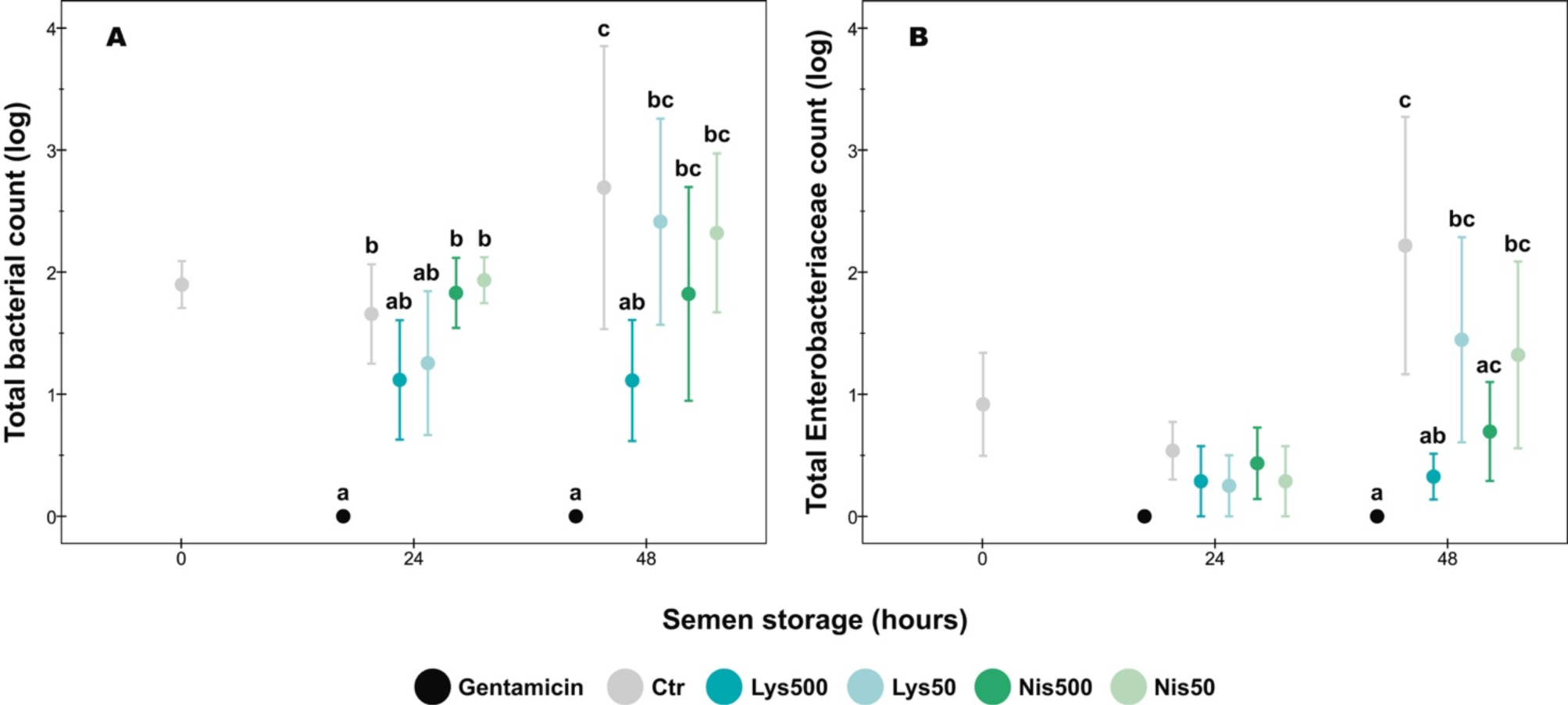
Effect of lysozyme and nisin on microbiological analysis of porcine semen during storage at 17 °C. A) Total bacterial count; B) Total Enterobacteriaceae count. Different superscript letters indicate significant differences (p<0.05) between treatments within each given time. The data are shown as the mean ± standard error of four replicates.

At 24 h of semen storage, the TBC were higher in Ctr and Nis treatments (p<0.05) than in the Gent treatment, while the latter did not differ from both Lys concentrations (p>0.05). At the same time, there were no significant differences in TEC between treatments (p>0.05).

At 48 h, only Lys500 did not differ from the Gent group in the TBC (p>0.05). Interestingly, there were no significant differences in TEC between the Gent group and Lys and Nis treatments at the highest concentration (500 µg/mL; p>0.05).

## Discussion

Our findings provide empirical evidence that both lysozyme and nisin enhance sperm parameters and reduce bacterial load during semen storage. Both AMPPs showed a higher percentage of motile sperm (at 24 h) and better-preserved sperm plasmalemma and acrosome integrity (24 h and 48 h) when compared to samples exposed to gentamicin. Moreover, lysozyme at 500 µg/mL did not show significant differences in the bacterial load (24 h and 48 h) and the percentage of sperm with rapid and progressive motility (SP1) compared to gentamicin treatment. On the other hand, Nisin at 500 µg/mL reduced the total number of Enterobacteriaceae but also decreased the percentage of sperm belonging to SP1 in comparison with the gentamicin group. The absence of toxicity of lysozyme to the sperm cells and its presence in the reproductive fluids of numerous animal species make this enzyme a suitable alternative to the common antibiotics used for boar semen preservation.

The use of AMPPs as alternative antibiotics is a promising approach for semen preservation as they are less likely to promote bacterial resistance because of their mechanism of action [30]. However, there are still challenges to cover such as their limited spectrum of antibacterial activity, noxious effects on sperm function, and their expensive and laborious production [30, 31]. Some AMPPs have been previously tested in boar semen showing a significant decrease in the bacterial load but also some toxicity to the sperm cells [32–34]. In the present study, lysozyme (500 µg/mL) kept the bacterial load at comparable levels to the samples treated with gentamicin without compromising the sperm function and even better preserving sperm acrosome and membrane integrity. In relation to sperm quality, the standard indicators established by breeding organizations worldwide for using preserved boar semen for AI are: 50-70% motile sperm and a bacterial load of <1000 CFU/mL [35]. Lysozyme at the highest concentration tested kept these parameters within the optimal range at 48 h of semen storage, with an averaged sperm motility of >65% and a bacterial load of <60 CFU/mL and 5 CFU/mL for total bacteria and Enterobacteriaceae counts, respectively. Nevertheless, the threshold on bacterial load in semen doses for AI is still widely debated. However, it is important to bear in mind that the different microbes usually detected in boar semen have also different toxicity to the sperm cells. For instance, bacteria such as *Alcaligenes* spp., *Actinomyces* spp., *Streptococcus* spp., and *Staphylococcus* spp. have almost no effects on sperm survival even in the presence of 10^10^–10^12^ CFU/mL; on the other hand, members of Enterobacteriaceae (i.e., *E. coli*, *Citrobacter* spp., *Klebsiella* spp., and *Serratia* spp.) together with *Proteus* spp. and *Pseudomonas* spp. have been classified as the most harmful bacteria to spermatozoa [5, 36]. For instance, *Pseudomonas* spp. can drop the seminal pH to 5.2–5.7 that results in a drastic decrease of sperm motility and acrosome integrity [36]. Even though using antibiotics, up to 32% of the semen doses are contaminated with several bacterial genera mainly because of AMR [4]. In this regard, Úbeda et al. [5] in a quality control of boar seminal doses (supplemented with antibiotics) established an above cut-off of 3×10^2^ CFU/mL for considering a semen sample as positive in bacterial contamination. On the other hand, some studies focusing on boar bacteriospermia [37] reported negative effects (litter size) when using semen for AI with more than 3.5×10^3^ CFU/mL (*E. coli*). According to these cut-offs for bacteriospermia, our findings show that Lys500 (in all replicates) is below the range that considers a sample as positive for bacterial contamination or the one that compromises sperm function and fertility outcomes. Although nisin treatments and Lys50 enhanced some sperm parameters when compared with Gent and Ctr groups, they had a TBC higher than the recommended range worldwide (<1000 CFU/ml) and they would be considered as positive for bacterial contamination (>300 CFU/mL; [5]).

The bacterial profile in boar semen closely depends on the hygienic conditions during the sample collection, the season, and the environmental characteristics where the animals are raised [38, 39]. In our work and like other studies [4] the most frequently isolated bacteria were *Pseudomonas aeruginosa* (G-), *Stenotrophomonas maltophilia* (G-), *Klebsiella* spp. (G-), and *Staphylococcus* spp. (G+). According to the abundance of *Staphylococcus* spp. observed in our study, it was reported that the presence of lysozyme in boar semen (2.4 µg/mL) might have been related to a bactericidal effect especially against *S. aureus* [40]. This finding, together with the role of lysozyme in the innate immunity and the great sperm-tolerance to this compound at high concentrations, indicates the suitability of this enzyme as an antimicrobial agent for boar semen preservation. The antimicrobial spectrum of lysozyme and nisin mainly includes G+ bacteria but, in combination with chelators, like EDTA, they can broaden their activity against G-. Semen extenders containing EDTA, such as the BTS, have been recently defined as “antimicrobially active extenders” as they allow to reduce the amount of antibiotic needed and act themselves as bacteriostatic in the absence of other antimicrobial agents [41]. Our results support these previous findings as both lysozyme and nisin (500 µg/mL) reduced the bacterial counts (Enterobacteriaceae) from 5.6×10^4^ CFU/mL (Ctr) to <14 CFU/mL (lysozyme: 5 CFU/mL; nisin: 13.75 CFU/mL). Both lysozyme and nisin have also shown the ability to reduce the endotoxic activity of LPS [42, 43], which is released by G-bacteria under antibiotic treatments (bacteriolysis) and negatively affects sperm quality. Gentamicin, on the other hand, cannot reduce the toxicity of LPS to the sperm cells at least at the concentration commonly used (250 µg/mL) for boar semen storage [30]. However, the combination of AMPPs with the common antibiotics used for sperm preservation neutralizes this bacteria-released endotoxin and increases sperm quality during semen storage [44]. The improved preservation of sperm membrane and acrosome integrity during semen storage, both for Lys and Nis treatments, could be therefore related to the capacity of these AMPPs to neutralise the detrimental effects of LPS on sperm function.

The presence of lysozyme in the semen of a wide range of invertebrate and vertebrate species is well known [45, 46]. In the seminal plasma, the abundance of this enzyme has been associated with good sperm quality [47, 48]. By contrast, human patients with chronic prostatitis have lower concentration of lysozyme than healthy men [49]. In spermatozoa, a lysozyme c-like protein (SLLP1) is located in the acrosome and involved in the fertilization process [50]. Similarly, the presence of a seminal vesicle-secreted lysozyme c-like protein (SVLLP) has been reported in mice. This protein binds to spermatozoa and suppresses bovine sperm albumin-induced sperm capacitation and inhibits acrosome reaction [51]. The enhanced sperm quality associated with an increased amount of lysozyme could be related not only to its antimicrobial properties but also to its ameliorative effect against oxidative stress. For instance, the oxidative damage caused to the sperm cells by the advanced glycation end-products (AGE), which is promoted by extenders containing high glucose concentration [52, 53], is cushioned by lysozyme activity [54]. It seems also plausible that this enzyme has no cytotoxic effects on sperm cells as gentamicin has [6, 7] because of its physiological presence in several body fluids including semen. On the other hand, nisin has shown spermicidal action (fast inhibition of sperm motility) in several mammalian species, including humans, in a range of concentrations from 50 to 400 µg/mL [55]. In our study, we did not observe such phenomena as we even found a significant enhancement in some sperm parameters (i.e., sperm motility −24 h- and acrosome/membrane integrity) compared to the gentamicin group. These differences could be attributed to nisin preparation (purification vs direct dilution in water), a high tolerance to this peptide in the porcine species, and reduced drug potency because of the presence of seminal plasma [55]. The enhancement of some sperm parameters by nisin treatments found in our study may be related to the recently reported antioxidant properties of this AMPP [56, 57]. This explanation is supported by the lower ORP values found at the highest concentration of nisin in the present study. However, in comparison to the gentamicin treatment, we also observed a decrease in some sperm velocity parameters and in the percentage of rapid and progressive spermatozoa (SP1), which might be explained by the impaired redox status [58] found in the samples treated with this AMPP. The bacterial load in nisin treatments (∼1700-3000 CFU/mL) on the second day could have also influenced the drop observed in kinetic parameters as the decline in sperm parameters due to bacterial contamination is more evident at 48 of semen storage [59, 60].

## Conclusions

Lysozyme (500 µg/mL) significantly reduces the bacterial load at comparable levels of samples treated with gentamicin (250 µg/mL). In addition, the sperm parameters (motility subpopulations, mitochondrial activity and, redox status) were unaltered or even better preserved (acrosome −24 h- and membrane integrity −48 h-) than in the gentamicin group. The presence of this enzyme in several body fluids (including semen and cervical mucus) and its sperm tolerance even at high concentrations, makes lysozyme an interesting alternative antimicrobial agent for boar semen preservation. Even though the bacterial load was low (<60 CFU/mL), our next steps are directed towards finding a natural compound that offers synergy against a broader spectrum of bacteria and the assessment of sperm fertilizing ability treated with this enzyme.

## Declarations

### Ethics approval and consent to participate

This study did not involve animal handling because the sperm samples were purchased as artificial insemination doses from a pig breeding company (Lipra Pork, Czech Republic).

### Authors’ contributions

JLR-S, PN, and EP: conceptualization; JLR-S: funding acquisition; JLR-S, PN, MS, and EP: methodology; JLR-S, MS, and EP: sperm analysis; PN: microbiological analysis; JLR-S: writing the first draft of the manuscript; PN, MS, and EP: review and editing. All authors contributed to manuscript revision, read, and approved the submitted version.

